# SpaCoEx: Sparse Gene Selection for Spatially Varying Co-expression in Spatial Transcriptomics

**DOI:** 10.64898/2026.09.05.749559

**Authors:** Michael Jian, Shaojun Pei, Gil Alterovitz

## Abstract

Spatial transcriptomics enables gene expression to be measured while preserving tissue location, but most existing analyses focus on spatial variation in individual genes or expression-defined domains. Here, we introduce SpaCoEx, a sparse spatial representation framework that integrates gene-expression levels with spatially varying gene-gene co-expression. SpaCoEx first performs structured gene selection then uses the selected genes to construct both expression-level features and local co-expression features. We applied SpaCoEx to two human spatial transcriptomics samples. In the cutaneous squamous cell carcinoma dataset, SpaCoEx selected 29 of 45 keratinocyte-related genes while preserving 96.74% of the spatial co-expression variation. In the breast cancer dataset, SpaCoEx identified spatially varying co-expression between B2M and HLA-C, a biologically meaningful major histocompatibility complex (MHC) class I antigen-presentation gene pair. Their local correlation was significantly higher in cancer-associated regions than in non-cancer regions (mean difference = 0.30, spatially adjusted SE = 0.045, *P* < 1.0 × 10^−10^), whereas B2M and HLA-C expression individually did not differ significantly between cancer and non-cancer. In benchmarking against manual tissue annotations, co-expression-only SpaCoEx achieved the strongest spatial coherence (percentage of abnormal spots [PAS] = 0.076), while the joint expression/co-expression representation achieved the highest annotation agreement, with an adjusted Rand Index (ARI) of 0.584 and normalized mutual information (NMI) of 0.663. By integrating marginal gene-expression information with local gene–gene co-expression structure, SpaCoEx provides a sparse, low-dimensional, and interpretable representation of spatial transcriptomics data that captures complementary aspects of tissue organization beyond expression-based variation alone.

## 1 Introduction

Spatial transcriptomics, which measures the expression level together with the spatial location of genes, has transformed the study of tissue organization [21, 11]. In contrast, single-cell RNA sequencing profiles gene activities at the cellular resolution but does not preserve cells’ original locations, and bulk RNA sequencing only provides averaged expression levels across diverse cell populations. By providing the physical locations of gene-expression measurements within intact tissue sections, spatial transcriptomics makes it possible to investigate how transcriptional programs are organized across anatomical structures, disease niches, and local microenvironments [4, 18].

Spatial context is especially important in heterogeneous tissues, where numerous distinct biological states can coexist within the same section. In breast cancer tissues, for example, normal epithelial areas, stromal compartments, immune-infiltrated niches, and transitional boundaries may occupy neighboring spatial domains while exhibiting distinct molecular profiles [26]. A major category of spatial transcriptomics analyses focuses on characterizing such spatial heterogeneity. Toward this goal, a number of analysis approaches have been proposed, such as identifying spatially variable genes and clustering spots into spatial domains [23, 30, 14, 16, 31, 28, 13]. For instance, SpatialDE detects genes whose expression level varies across spatial locations [23], and SPARK quantifies spatial expression patterns through the use of generalized linear spatial models [22]. Although these approaches have offered important views of tissue heterogeneity, they primarily describe spatial variation in individual genes or cell-type composition.

Biological functions are rarely determined by individual genes acting in isolation. Genes work through coordinated regulatory programs, signaling pathways, and cell-state-dependent networks. Early gene-expression studies found that genes with similar expression patterns can be grouped into biologically meaningful clusters [10]. Co-expression network methods such as Weighted Gene Coexpression Network Analysis have demonstrated that gene-gene relationships often reveal functional modules and network structure [29, 25]. Two tissue regions may show similar expression levels for a set of genes but differ substantially in their correlation structures. Specifically, a pair of genes that are strongly co-expressed in one spatial region might be weakly co-expressed in another, which would be due to local distinctions in cell composition, regulatory state, tumor progression, tissue structure, or micro-environmental signaling. Thus, spatial tissue organization may be reflected not only in variation in gene expression levels, but also in variation in gene-gene co-expression relationships.

Several recent methods have been proposed to understand spatially varying gene co-expression networks. For example, SpatialCorr tests whether the correlation structure of a predefined gene set varies spatially [5], and SpaceX estimates gene co-expression networks in spatially structured tissues by accounting for spatial correlation and tissue domains [2]. More recent approaches further model spatially informed gene co-expression networks by jointly capturing spatial dependence across locations and co-expression among genes [24]. While these developments suggest that spatial transcriptomics analysis provides additional biological information over marginal gene-level spatial variation, they often require a given set of genes to be analyzed or use gene screening methods that are not designed to identify spatially varying gene co-expression. This is because, as the number of genes increases, local co-expression matrices grow quadratically in dimension, and many genes may contribute redundant or noisy information. Including all genes can obscure the dominant spatial structure of co-expression and reduce the interpretability of downstream analyses. Therefore, a central problem is to select a sparse but informative subset of genes that preserves the major spatial variation in local co-expression structure.

Here, we introduce SpaCoEx, a framework for constructing sparse spatial representations that integrate both gene-expression levels and spatially varying gene-gene co-expression. SpaCoEx first computes local co-expression matrices at each spatial location using neighboring spots and applies a log-co-expression transformation to enable principled comparison of co-expression structures across tissue space. It then formulates gene selection as a structured optimization problem, in which selecting or removing a gene corresponds to retaining or excluding the full row and column associated with that gene in the local co-expression matrices. By balancing preservation of spatial co-expression variation with sparsity, SpaCoEx identifies genes that contribute most strongly to spatially varying co-expression.

Using the selected genes, SpaCoEx constructs two complementary feature representations: expression features, which capture the abundance of individual genes across spatial locations, and co-expression features, which describe how relationships among these genes vary across local tissue neighborhoods. These two representations are then combined through a weighted joint representation, where the parameter *α* controls the relative contribution of expression and co-expression information. This formulation allows SpaCoEx to compare expression-only, co-expression-only, and joint representations within the same framework, and to evaluate whether integrating marginal expression levels with local gene-gene relationship structure improves downstream analyses, including dimensionality reduction, spatial clustering, and detection of biologically meaningful tissue domains such as cancer-enriched regions.

## 2 Materials and methods

### 2.1 Overview of the SpaCoEx framework

SpaCoEx is designed to extract a sparse representation of spatially varying co-expression from spatial transcriptomics data. A visual representation of the workflow can be seen in Figure 1. Given a spatial transcriptomics dataset with gene expression measurements, along with spatial coordinates, SpaCoEx first computes the local gene-gene co-expression (correlation) matrix for each spot by using its nearest neighbors. In this step, a matrix logarithm maps correlation matrices into a log-Euclidean representation. SpaCoEx then performs a structured *L*_1_-penalized gene selection procedure to identify genes that preserve the most spatial variation in local co-expression matrices. Note that gene selection is structured at the level of whole rows and columns, rather than individual matrix entries, because selecting a gene keeps all matrix entries involving that gene.

**Figure 1.**
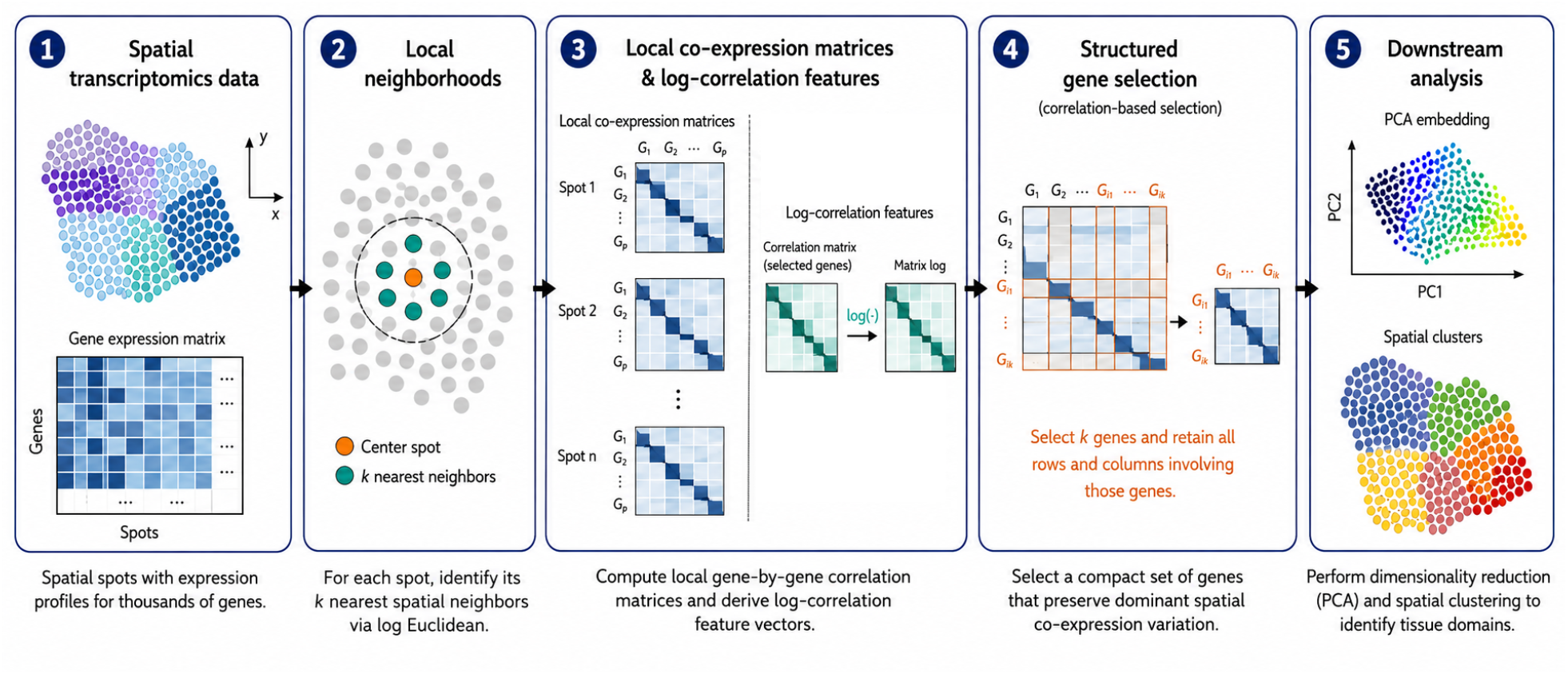
Overview of the SpaCoEx Workflow.

Based on the selected genes, SpaCoEx constructs a local co-expression representation and applies principal component analysis (PCA) to summarize the dominant axes of spatial co-expression variation. The resulting PC features are used later for downstream analysis. To evaluate the usefulness of co-expression features, we also propose a fused approach to combine expression-level and co-expression features in a weighted manner.

### 2.2 Local neighborhood construction

For each spatial spot, SpaCoEx defines a local neighborhood using the spatial coordinates of the tissue section. Let *s* = 1, …, *n* index the spatial spots. For a fixed neighborhood size *K*, the local neighborhood *N*_*K*_(*s*) is defined as the set of *K* spatial spots closest to spot *s* in physical tissue space, including spot *s* itself.

For each spot *s*, the local co-expression *C*(*s*) ∈ ℝ^*p×p*^ is defined as the correlation matrix of the genes using the expression values from spots in N_*K*_(*s*), where *p* is the number of genes. The (*i, j*)th entry of *C*(*s*) represents the local correlation between genes *i* and *j* around spot *s*. Thus, SpaCoEx converts the spatial transcriptomics dataset into a collection of local co-expression matrices,

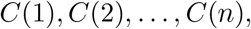

one for each spatial spot.

The neighborhood size *K* controls the spatial scale at which local co-expression is estimated. Smaller values of *K* use fewer nearby spots and may capture more local variation, while larger values of *K* average over broader tissue regions. In our analysis, SpaCoEx uses *K* = 15 nearest neighbors.

### 2.3 Log-Euclidean Representation of Local Co-expression Matrices

Unlike individual gene expression features, local co-expression matrices do not form a standard Euclidean vector space. A local correlation matrix is naturally symmetric positive semi-definite. To ensure positive definiteness, we add a small positive constant to obtain a symmetric positive definite (SPD) matrix. The space of SPD matrices possesses a curved geometric structure, forming a Riemannian manifold rather than a flat vector space. Accounting for this geometry is crucial because standard Euclidean operations such as direct matrix addition or subtraction do not preserve the underlying manifold structure. To consistently compare local co-expression matrices across different spatial spots, we adopt a log-Euclidean framework [3]. The matrix logarithm maps each SPD co-expression matrix from its curved manifold into the vector space of symmetric matrices. Within this transformed space, matrix differences can be naturally and rigorously evaluated using the Frobenius norm.

For each spot *s*, we denote the local correlation matrix by *C*(*s*). In practice, we use the neighboring spots of *s* to compute it. The log of a correlation matrix is defined using the eigenvectors and logtransformed eigenvalues. Because the matrix logarithm is only well-defined for positive definite matrices, SpaCoEx first regularizes each local correlation matrix by adding a small ridge term:

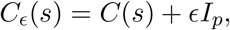

where *I*_*p*_ is the *p* × *p* identity matrix and *ϵ >* 0 is a small ridge constant.

Let 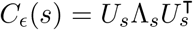 be the eigendecomposition of the regularized local correlation matrix. Then, the log co-expression matrix is defined as

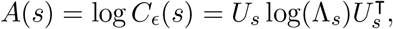

where log(Λ_*s*_) is obtained by applying the logarithm to each eigenvalue. After applying the matrix logarithm, each local co-expression matrix is represented as a symmetric matrix *A*(*s*). These log co-expression matrices are used for structured gene selection. After gene selection, the upper triangular entries of the selected log co-expression matrices are vectorized to form SpaCoEx features for dimensionality reduction and clustering.

### 2.4 Structured gene selection objective with *L*_1_ penalty

After constructing local log co-expression matrices, the next step is to select a subset of genes while preserving the important spatial variation in co-expression.

Let

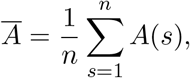

denote the average co-expression across all spots. Note that the centered matrix *A*(*s*) − *Ā* describes how the local co-expression around a spot *s* differs from the average co-expression structure across the tissue. The gene selection problem aims to maximize the variation of co-expression across spots, which is defined as 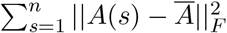.

Note that this is not a standard feature-selection problem, because here, the features are organized as matrices. This means that removing a gene should remove all the effects of the gene, including co-expression entries. For example, as shown in Figure 2, the matrix entries associated with gene *j* are

**Figure 2.**
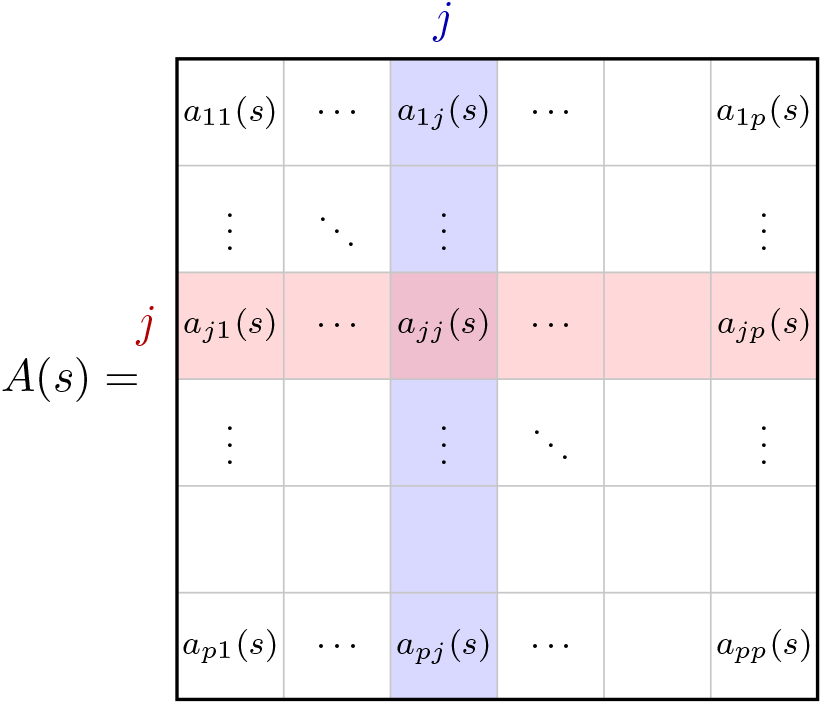
Gene selection is structured because gene *j* corresponds to all entries in row *j* and column *j*.

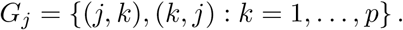

This means that gene selection here must be an all-in or all-out situation. If gene *j* is excluded, then all the spatial variation is not expressed, and vice versa. As such, this method relies on a group-structured selection idea.

To enforce gene-level selection, we introduce a continuous weight *w*_*j*_ ∈ [0, 1] to gene *j*. Let *W* denote the diagonal matrix of the weights, then the (*i, j*)th entry of (*A*(*s*) − *Ā*)*W* is

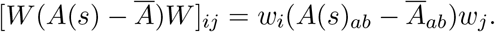

Thus, the co-expression between genes *i* and *j* is retained if and only if when *w*_*i*_*w*_*j*_ *>* 0. On the other hand, by setting *w*_*j*_ to be 0, every entry involving gene *j* also becomes 0, since both the row and column weight is 0. Thus, row and column *j* are removed.

To select a sparse set of genes that can preserve as much variation in the local log co-expression matrices as possible, we introduce an *L*_1_ penalty to encourage some of the weights to be zero. Combining the preservation term and the sparsity penalty yields the final optimization problem,

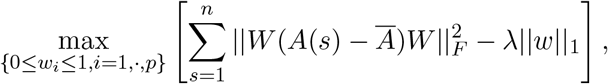

where *λ* denote the amount of penalty.

### 2.5 A Projected Proximal Gradient Ascent Algorithm

The objective can be written as

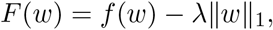

where 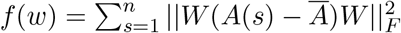 is the smooth part of *F* (*w*).

As derived in the Supplementary data, the projected proximal gradient ascent algorithm consists of the following three steps.

- First, take a gradient ascent step on the smooth preservation term:

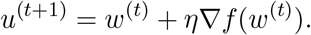
- Second, apply the proximal step for the *L*_1_ penalty. Because the weights are constrained to be nonnegative, this becomes a one-sided soft-thresholding step:

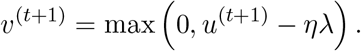

This step shrinks each weight by an amount proportional to *λ*. If a weight becomes small enough, it is set exactly to zero, which removes the corresponding gene.
- Third, project the weights back into the allowed interval [0, 1]:

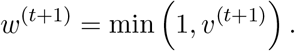

Equivalently, the full update can be written as

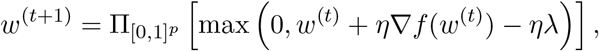

where 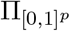 denotes projection onto the box [0, 1]^*p*^.

### 2.6 Parameter Tuning

The sparsity parameter *λ* controls the tradeoff between preserving spatial variation and selecting fewer genes. To study this tradeoff, we run the optimization over a grid of *λ* values from 10^−1^ to 10^0^. For each value of *λ*, we compute the optimized weight vector *w*_*λ*_ and the corresponding diagonal matrix *W*_*λ*_, and the preserved variation,

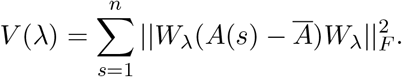

Thus, the preserved fraction is

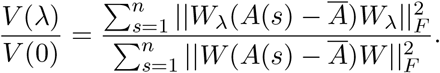

This preserved fraction provides an interpretable way to compare different choices of *λ*. A value close to 1 means that the selected genes retain most of the spatial variation in the local co-expression matrices, while a smaller value means that more variation has been lost. By comparing the preserved fraction with the number of selected genes, a sparsity level can be chosen that reduces the number of selected genes, while still keeping most of the co-expression signal.

### 2.7 Joint expression and co-expression representation

The expression and co-expression characterize, respectively, how individual genes and gene-gene coordination vary across spatial locations. To combine information from them, we propose to conduct PCA and then choose the top 20 principal components from each to represent underlying features.

Let *E* ∈ ℝ^*n×*20^ denote the expression PCA scores and let *C* ∈ ℝ^*n×*20^ denote the co-expression PCA scores. Because the two representations can have different overall magnitudes, each score matrix was normalized using its Frobenius norm:

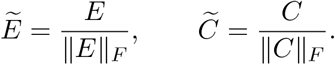

For an expression weight *α* ∈ [0, 1], the joint representation was defined as

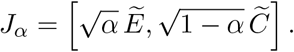

Then the squared distance between spots *i* and *j* in this representation is

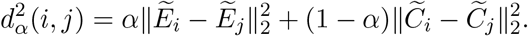

Thus, *α* directly controls the relative contribution of expression to the distance used for clustering. When *α* = 0, the representation is co-expression-only; when *α* = 1, it is expression-only; and intermediate values combine both sources. We evaluated *α* ∈ {0, 0.25, 0.50, 0.75, 1} and applied k-means with ten clusters to each representation.

### 2.8 Datasets

We applied SpaCoEx to two publicly available spatial transcriptomics samples. The first sample was originally generated by Ji et al. [15] as part of a multimodal study of composition and spatial architecture in human cutaneous squamous cell carcinoma, and is available through the Gene Expression Omnibus. In this report, we analyzed the spatial transcriptomics sample `GSM4284326`, also referred to as `P10_ST_rep2`. The sample contains gene expression measurements together with spatial coordinates for 621 spots across the tissue section. After preprocessing, the dataset contained 16,642 genes. In this analysis, SpaCoEx was applied to the subset of 45 keratinocyte-related genes, because keratinocyte programs are biologically relevant to squamous cell carcinoma tissue organization.

We then assessed SpaCoEx using an annotated breast cancer dataset. This dataset is commonly known as Human Breast Cancer Block A Section 1 [1]. Manual annotations for the breast cancer sample were obtained from the `BRCA1` metadata file in the SEDR analyses repository [27], which provides both coarse labels and finer region labels for each spatial barcode. These annotations group spots into healthy tissue, tumor, invasive carcinoma, and surrounding tumor regions, with finer labels corresponding to IDC, DCIS/LCIS, tumor-edge, and healthy subregions.

## 3 Results

### 3.1 SpaCoEx identifies spatially varying co-expression in cutaneous squamous cell carcinoma regions

Recall that for the cutaneous squamous cell carcinoma sample we focus on the 45 keratinocyte-related genes. For each spatial spot, local co-expression was estimated using the 15 nearest neighboring spots to calculate a correlation matrix for each spot. To choose the number of genes, we examined the tradeoff between sparsity and the preserved spatial co-expression variation, through the use of a *L*_1_ penalty. As the penalty *λ* increases, fewer genes are selected, the preserved fraction decreases, as shown in Figure 3 B. The goal is to identify a smaller subset of genes that retains most of the spatially varying co-expression signals. In this analysis, SpaCoEx selected 29 of the 45 keratinocyte genes, preserving a *>* 90% of the spatial co-expression variation among the original keratinocyte gene set (see Figure 3 A and B).

**Figure 3.**
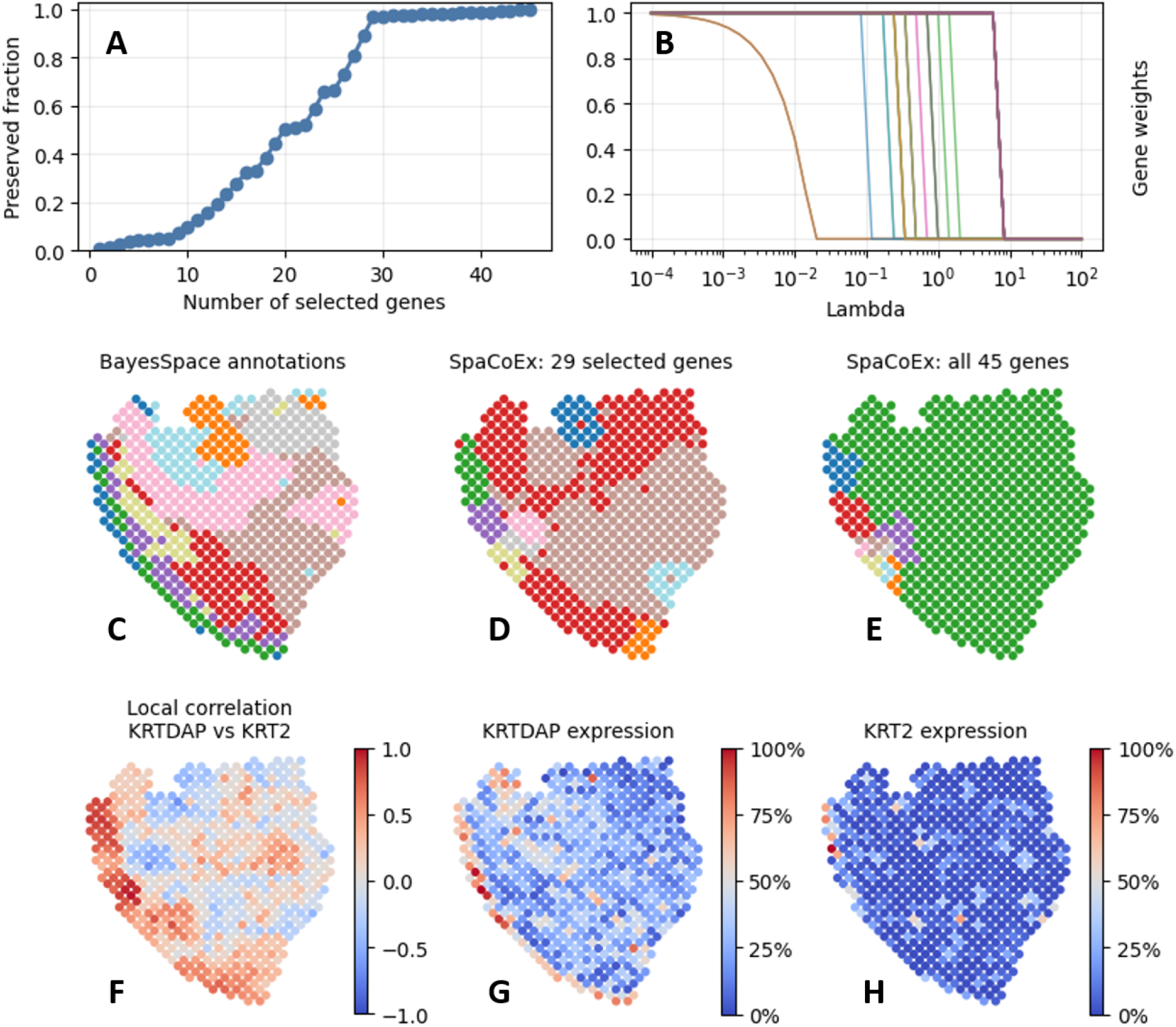
SpaCoEx results for the cutaneous squamous cell carcinoma sample. A) the preserved fraction of spatial co-expression variation over number of genes selected. SpaCoEx selected 29 genes, preserving *>* 90% of the spatial co-expression variation among the original keratinocyte gene set. B) The gene weight path as *λ* increases. C) clusters using BayesSpace. D) SpaCoEx clusters using the 29 selected keratinocyte genes. E) SpaCoEx clusters using all 45 keratinocyte genes. F) local correlation between *KRTDAP* and *KRT2* across the tissue. G) spatial expression of *KRTDAP*. H) spatial expression of *KRT2*. These plots illustrate how local co-expression can reveal spatial variation in gene-gene relationships that is not fully captured by individual gene expression maps.

We then examine the clusters based on the 29 selected genes. To do that, we first vectorized the log-transformed correlation matrices using the 29 genes, computed the top 20 principal components, and then performed k-means with 10 clusters. Figure 3 D shows that the co-expression features based on the 29 selected genes produced clustering with reasonable similarity to that of BayesSpace (ARI=0.15, NMI=0.28), whereas the all-gene analysis appeared to miss some clusters, potentially because irrelevant genes added noise.

To examine what the selected genes represent at the local co-expression level, we visualized the first two genes selected by SpaCoEx, *KRTDAP* and *KRT2*, in Figure 3 F-H. Both genes are related to keratinocyte and epidermal differentiation: *KRTDAP* has been described as a keratinocyte-associated gene involved in epithelial stratification, while *KRT2* is an epidermal keratin expressed during later stages of keratinocyte differentiation [19, 7]. Therefore, changes in their local relationship across the tissue may reflect spatial differences in keratinocyte differentiation state or tissue organization.

As shown in Figure 3 F, the local correlation between *KRTDAP* and *KRT2* varies across the tissue section. This means that even when two genes have spatially structured expression patterns, the strength and direction of their co-expression can still change across tissue regions. This example illustrates spatial variation in gene-gene relationships that may not be apparent from individual gene expression levels.

Interestingly, Guo et al. [12] suggest that *KRT2* and *KRTDAP* may co-vary because both are associated with the same spatially informed component. Our result is consistent with known gene- or protein-level associations involving *KRT* -family genes. This suggests that our gene selection method can identify genes whose co-expression patterns vary spatially.

This gene-selection step is important because local co-expression matrices grow proportional to the square of the number of genes. Reducing the gene set decreases the dimension of the local co-expression representation and improves interpretability, while the preserved-fraction curve provides a direct way to quantify how much spatial co-expression variation is retained. Perhaps more important, the identified spatially varying co-expression, such as the *KRT2* and *KRTDAP* pair, might provide useful biological insights for understanding how genes coordinate in different tissue regions. This example with *KRT* family genes serves as a demonstration of SpaCoEx. Although it also shows satisfactory clustering performance relative to existing methods, we focus on the method comparison on a manually annotated breast cancer sample.

### 3.2 SpaCoEx identifies spatially varying co-expression in cutaneous squamous cell carcinoma regions

#### 3.2.1 SpCoEx identified spatially varying gene-gene co-expression and differential co-expression between cancer and non-cancer regions

For the Human Breast Cancer Block A Section 1 sample, we first retrained genes in at least 50% cells and selected top 200 based on Moran’s index. SpaCoEx was then applied to this candidate gene set and keep the 139 genes that preserve 50% of variance in co-expression. Note that we used a 90% variance threshold in the previous example because there is a clear elbow in the variation plot and the analysis focused on a predefined set of candidate genes (i.e., the *KRT* family), whereas here there is no obvious elbow in the explained variation plot and the candidate genes had already been broadly pre-selected based on expression abundance and spatial variation. Therefore, consistent with common practice in PCA-based analyses of gene expression [17], we considered retaining 50% of the co-expression variance to be a more appropriate threshold in this setting.

Among the top ranked genes selected by SpaCoEx, many showed spatially varying co-expression with other genes. For example, Figure 4 shows that the correlation between *B2M* and *HLA-C* changes substantially across spatial locations. It is also notable that the local correlation map is largely consistent with cancer-associated regions, suggesting that this co-expression relationship is spatially associated with cancer-enriched tissue. Note that the expression maps of either gene alone do not align as closely with these regions.

**Figure 4.**
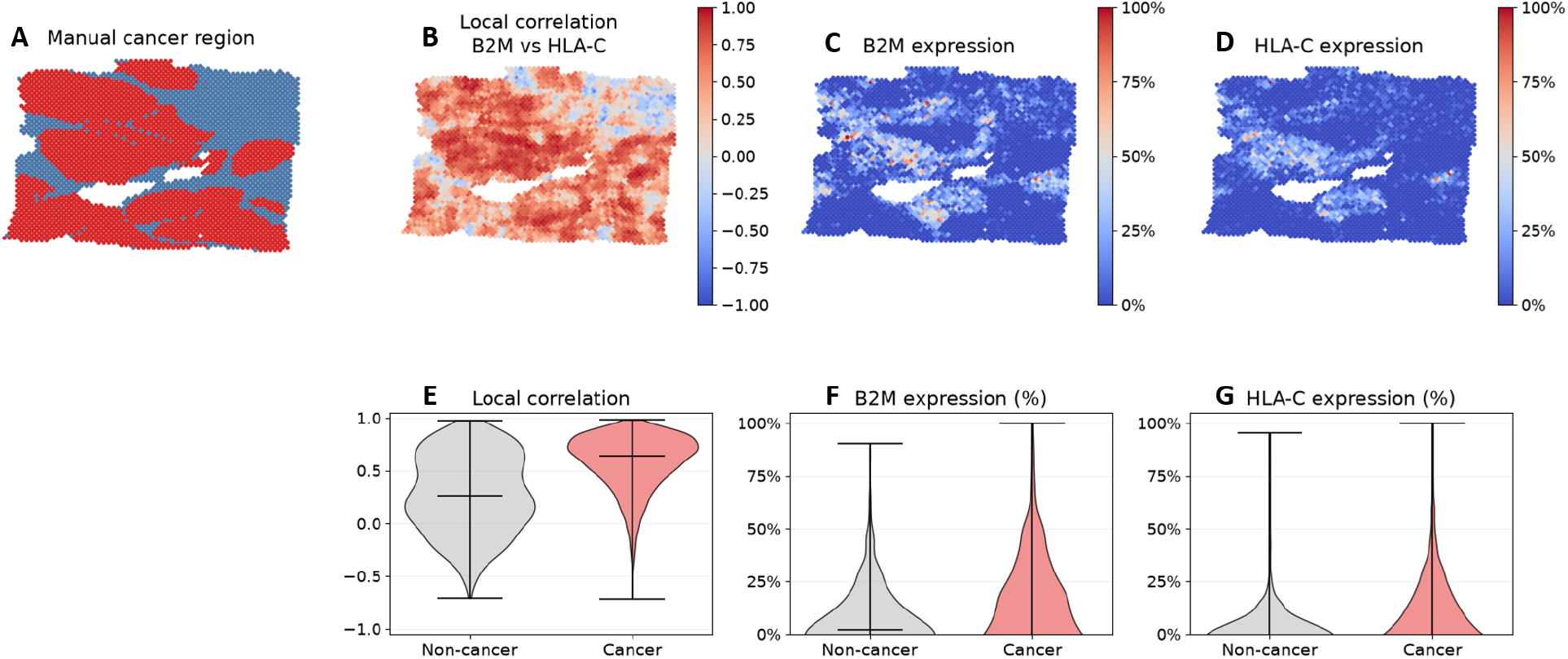
Spatial relationship between *B2M* and *HLA-C* in the breast cancer dataset. A) manually annotated cancer region. B) local correlation between *B2M* and *HLA-C* across the tissue. C) spatial expression of *B2M*. D) spatial expression of *HLA-C*. The local correlation map more clearly follows the cancer-associated spatial structure than either expression map alone. E) violin plots of local correlations in Cancer and non-cancer regions. F) violin plots of *B2M* expression. G) violin plots of *HLA-C* expression.

We then estimated cancer vs non-cancer difference using the observed difference in mean expression / correlation. To account for spatial dependence among neighboring spots, standard errors, confidence intervals, and P values were calculated using Conley spatial heteroskedasticity- and autocorrelation-consistent standard errors [8], implemented in the R package `conleyreg` [9]. Euclidean distances between spot coordinates were used with a Bartlett kernel and a cutoff of 2,000 spatial units, selected based on the residual semivariogram. Other reasonable choices of the cutoff lead to similar results in significance levels.

Consistent with Figure 4, the difference between cancer and non-cancer is significant for local correlation (mean difference = 0.30, spatially adjusted SE = 0.045, *P <* 1.0 × 10^−10^), but not for *B2M* expression (mean difference = -0.48, spatially adjusted SE = 0.33, *P* = 0.15) or for *HLA-C* expression (mean difference = 0.04, spatially adjusted SE = 0.26, 0.87). This indicates that the relationship between *B2M* and *HLA-C* captures spatial information relevant to cancer-associated tissue structure beyond either gene’s expression pattern alone. *B2M* encodes beta-2-microglobulin, which is required for stable *MHC* class I expression, whereas *HLA-C* encodes a classical *MHC* class I heavy chain. Their coordinated expression is therefore important for *MHC* class I antigen presentation. Consequently, the spatially varying local correlation between these genes may reflect regional differences in antigen presentation and the immune landscape of the tumor microenvironment.

#### 3.2.2 Comparison with existing spatial transcriptomics methods

Unlike the cutaneous squamous cell carcinoma analysis, this dataset includes manual tissue annotations (Figure 5), which makes it possible to compare clustering results with known tissue regions. We therefore compare the clustering results of SpaCoEx and several other methods, including Scanpy Leiden, stLearn, and GraphST. These methods use expression profiles, spatial coordinates, graph structure, and, for some methods, histology to identify spatial domains. SpaCoEx instead creates local gene-gene co-expression features and then clusters spots based on variation in local co-expression structure. Thus, SpaCoEx provides a complementary view of tissue organization; it groups spots not only by the expression levels of genes, but also by how the genes vary together locally.

**Figure 5.**
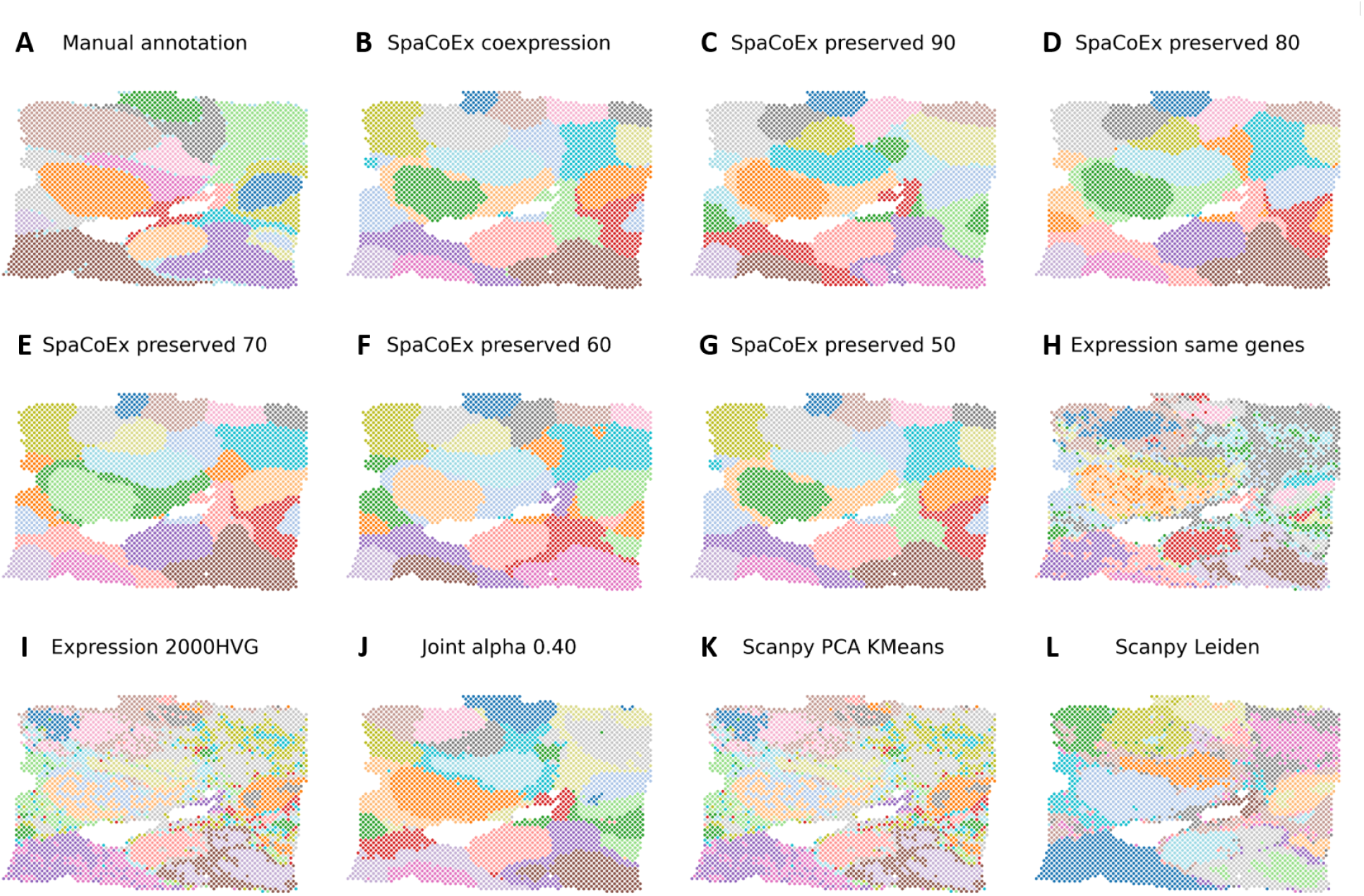
Spatial domain comparison on annotated breast cancer spatial transcriptomics data. A) the manual annotation is shown alongside SpaCoEx co-expression (B-G), expression-only (H, I), joint expression/co-expression (J), and existing spatial transcriptomics methods (K, L).

**Table 1.** Conceptual comparison between SpaCoEx and related spatial transcriptomics methods.

| Method | Primary goal | Difference from SpaCoEx |
| --- | --- | --- |
| Scanpy Leiden | Graph-based clustering of cells/spots using expression similarity | SpaCoEx clusters using spatially local co-expression structure, whereas Leiden typically operates on a neighborhood graph derived from expression profiles. |
| stLearn | Spatial analysis using expression, spatial location, and histology | SpaCoEx focuses on local gene-gene relationship structure. |
| GraphST | Graph representation learning for spatial transcriptomics | SpaCoEx constructs interpretable co-expression features instead of learned graph embeddings. |

To evaluate the spatial-domain results quantitatively, we used three complementary metrics. The adjusted Rand index (ARI) and normalized mutual information (NMI) measure agreement with the manual annotations, with ARI additionally correcting for chance agreement. The percentage of abnormal spots (PAS) measures spatial label discontinuity; lower values indicate greater spatial coherence. Together, ARI and NMI evaluate concordance with the reference annotations, whereas PAS provides a complementary assessment of the spatial coherence of the inferred domains.

We compared SpaCoEx co-expression features, expression-only features, joint expression and co-expression features, and several existing spatial transcriptomics methods. For existing methods, we used the top 200 genes based on Moran’s index and their default parameters. 10 clusters was assumed in all methods. Clustering agreement with the manual annotation was measured using ARI and NMI, while spatial coherence was measured using PAS. Lower PAS values indicate that neighboring spots are more often assigned to the same cluster.

**Table 2.** Benchmarking SpaCoEx and comparison methods on annotated breast cancer spatial transcriptomics data. ARI and NMI compare agreement with the manual annotation, while PAS measures spatial label discontinuity.

| Method | ARI | NMI | PAS |
| --- | --- | --- | --- |
| SpaCoEx co-expression only | 0.428 | 0.641 | **0.076** |
| Expression only, same genes | 0.408 | 0.565 | 0.426 |
| Expression only, 2000 HVGs | 0.395 | 0.486 | 0.495 |
| Joint, $\alpha = 0.40$ | 0.534 | **0.663** | 0.116 |
| Joint, $\alpha = 0.60$ | **0.584** | 0.641 | 0.235 |
| stLearn | 0.499 | 0.572 | 0.337 |
| Scanpy Leiden | 0.424 | 0.568 | 0.366 |
| GraphST | 0.352 | 0.497 | 0.468 |

The breast cancer benchmark shows that co-expression contains a strong spatial signal. SpaCoEx co-expression features produced the lowest PAS value, indicating the most spatially coherent clustering. However, the strongest agreement with the manual annotation was obtained by the joint representation. In particular, *α* = 0.60 produced the highest ARI, while *α* = 0.40 produced the highest NMI.

These results suggest that expression and co-expression capture complementary aspects of tissue organization. Expression information improves agreement with manually labeled tissue regions, while co-expression information improves spatial coherence. Compared with expression-only baselines and existing spatial transcriptomics methods, the joint SpaCoEx representation produced stronger agreement with the manual annotation.

## 4 Discussion

In this study, we developed SpaCoEx as a sparse spatial representation framework for integrating gene-expression levels with spatially varying gene-gene co-expression in spatial transcriptomics data. Most existing spatial transcriptomics analyses characterize tissue organization through spatially variable genes, expression-based clustering, or the integration of expression with spatial coordinates and histology [21, 23, 22, 30, 14, 16]. SpaCoEx addresses a complementary question: whether local changes in gene-gene relationships can reveal spatial tissue structure beyond marginal expression levels alone. By combining expression features with local co-expression features, SpaCoEx provides a joint representation of both individual gene abundance and local gene coordination.

A key methodological contribution of SpaCoEx is that it reframes gene selection for spatial transcriptomics as a structure-preserving selection problem on a field of local co-expression matrices. Existing gene-selection strategies typically prioritize genes according to marginal expression variation or spatial expression patterns, but these criteria do not directly evaluate whether a gene contributes to spatial changes in gene-gene relationship structure. SpaCoEx instead defines gene importance by how much the full set of co-expression entries associated with that gene contributes to variation in local log-co-expression matrices across tissue space. Because each gene corresponds to an entire row and column in the co-expression matrix, the method selects or removes all pairwise relationships involving that gene as a single structured unit. In this way, SpaCoEx extends the idea of group-structured selection to spatially varying matrix-valued co-expression data, producing a sparse gene set that preserves the dominant spatial co-expression signal before downstream feature construction, dimensionality reduction, and clustering. The log-Euclidean representation further enables these local matrix-valued features to be compared in a principled Euclidean space [3]. This structured selection also provides an important dimensional advantage. For *p* genes, the vectorized upper triangular entries of a local co-expression matrix contain *p*(*p* + 1)*/*2 features before downstream dimension reduction. Thus, the feature dimension grows quadratically with the number of genes. By selecting genes before constructing the final co-expression representation, SpaCoEx reduces the dimensionality of the problem while retaining interpretable gene-level structure and produces a lower-dimensional and more interpretable representation without ignoring the matrix structure of gene-gene relationships.

The biological examples support the value of modeling local co-expression. In the cutaneous squamous cell carcinoma dataset, SpaCoEx identified spatially varying co-expression between *KRTDAP* and *KRT2*, two genes related to keratinocyte and epidermal differentiation [7, 19]. The local relationship between these genes varied across the tissue section, suggesting that SpaCoEx can capture regional differences in keratinocyte differentiation or epithelial organization through gene coordination rather than through expression levels alone. In the breast cancer dataset, SpaCoEx identified *B2M* and *HLA-C* as a spatially varying gene pair associated with cancer-enriched regions. This pair is biologically meaningful because *B2M* encodes beta-2-microglobulin, which is required for stable *MHC* class I expression, whereas *HLA-C* encodes a classical *MHC* class I heavy chain involved in antigen presentation [6, 20]. The stronger local correlation between *B2M* and *HLA-C* in cancer-associated regions suggests that spatially varying co-expression may reflect regional differences in antigen presentation and tumor-immune organization.

Compared with existing spatial-domain identification methods, SpaCoEx showed strong performance in the annotated breast cancer benchmark. The joint expression/co-expression representation achieved stronger agreement with manual tissue annotations than expression-only clustering and the tested spatial transcriptomics methods, including Scanpy Leiden, stLearn, and GraphST. In particular, the joint representation achieved the highest ARI, whereas the co-expression-only representation achieved the lowest PAS, indicating the strongest spatial coherence. These results suggest that the advantage of SpaCoEx does not come from co-expression alone, but from explicitly modeling two complementary sources of spatial information: marginal gene-expression variation and local gene-gene coordination. Expression features improve alignment with manually annotated tissue regions, whereas co-expression features preserve spatially coherent local relationship structure. Thus, SpaCoEx provides a joint spatial representation that improves spatial-domain characterization while remaining interpretable at both the gene-expression and gene-pair relationship levels.

Several limitations remain. First, SpaCoEx has currently been evaluated on a limited number of datasets, and additional studies across tissue types, disease contexts, and platforms are needed to assess generalizability. Second, the selected genes depend on the initial candidate gene set, making robust gene pre-filtering an important practical step. Third, local co-expression estimation depends on the neighborhood size, which determines the spatial scale of the inferred gene-gene relationships. Future work should evaluate adaptive or Gaussian-weighted neighborhoods and assess the stability of selected genes, local co-expression patterns, and spatial domains across spatial scales. Finally, SpaCoEx is currently applied to one tissue section at a time. Extending the framework to multiple samples may allow spatially varying expression and co-expression features to be used for predicting disease states, aging, tumor progression, or treatment response across cohorts.

Overall, SpaCoEx provides a sparse, low-dimensional, and interpretable framework for studying spatial transcriptomics by integrating marginal gene-expression variation with local gene-gene coexpression structure. These results suggest that spatial tissue heterogeneity is reflected not only in where genes are expressed, but also in how genes coordinate with one another across local tissue neighborhoods.

### Key Points

- We proposed SpaCoEx, a computational framework to identify genes showing spatially varying co-expression patterns.
- *L*_1_ regularization was used to encourage sparsity in the set of selected genes.
- SpaCoEx identified genes that show differential correlation between breast cancer and noncancer regions but are not marginally differentially expressed between the two types of regions.

## 5 Biographical Note

Michael Jian is a participant of the 2026 MIT PRIMES-USA program. Dr. Shaojun Pei is a postdoctoral researcher in the Division of General Internal Medicine and Primary Care at Brigham and Women’s Hospital. Dr. Gil Alterovitz is an associate professor at Harvard Medical School.

## 8 Data and code availability

Both datasets presented in this article are openly available. Links to the data are provided below.

Human cutaneous squamous cell carcinoma (accessed on 2026-6-12): https://www.ncbi.nlm.nih.gov/geo/query/acc.cgi?acc=GSM4284326

Human Breast Cancer Block A Section 1 (Accessed on 2026-08-08) https://www.10xgenomics.com/datasets/human-breast-cancer-block-a-section-1-1-standard-1-0-0

The code used in this paper: https://github.com/mcjian09/SpaCoEx

## 9 Author contributions

Michael Jian designed the research, developed software, and wrote the manuscript; Shaojun Pei designed the research and wrote the manuscript; Gil Alterovitz designed the research and wrote the manuscript.

## 10 Supplementary data

### 10.1 A Projected Proximal Gradient Ascent for Gene Selection in Spatially Varying Co-Expression

Let 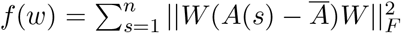 . For a given amount of penalty *λ >* 0, the objective can be written as

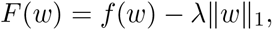

and the goal is to maximize *F* (*w*) subject to 0 ≤ *w*_*i*_ ≤ 1, *i* = 1, ·, *p*. Because all weights are non-negative,

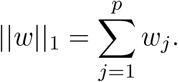

The parameter *λ* controls the strength of the penalty.

Define

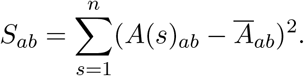

The matrix *S* records how much each gene-gene log-correlation entry varies across space. Larger values indicate greater spatial variation in the corresponding log-correlation entry.

It follows that

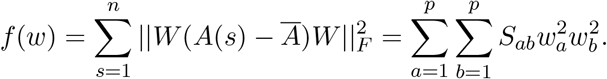

Substituting these expressions, we can rewrite the optimization problem as

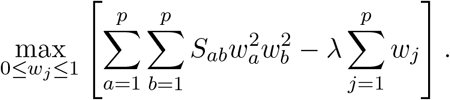

Ordinary gradient ascent is not sufficient for this maximization problem because the objective contains both a non-smooth penalty and constraints on the weights. The preservation term *f* (*w*) is smooth and differentiable, so it can be handled using a gradient ascent step. However, the *L*_1_ penalty is not differentiable when *w*_*j*_ = 0. This matters because exact zeros are desired: when *w*_*j*_ = 0, gene *j* is removed from the selected set. A standard gradient method would shrink weights continuously, but would not naturally create exact zeros. In addition, the weights must satisfy 0 ≤ *w*_*j*_ ≤ 1, since each *w*_*j*_ represents the selection strength of gene *j*. An unconstrained gradient step could move weights below 0 or above 1, which would make them invalid. Therefore, we use projected proximal gradient ascent: the gradient step increases the smooth preservation term, the proximal step handles the *L*_1_ penalty, and the projection step keeps the weights in the valid interval.

The gradient of the smooth term is derived as follows. Since

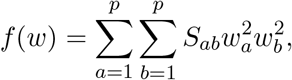

the derivative with respect to *w*_*j*_ is

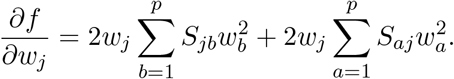

Since *S* is symmetric, this becomes

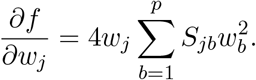

In vector form,

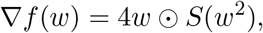

where ⊙ denotes elementwise multiplication, and *w*^2^ means that each entry of *w* is squared.

The projected proximal gradient ascent update has three steps. First, take a gradient ascent step on the smooth preservation term:

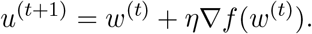

Second, apply the proximal step for the *L*_1_ penalty. Because the weights are constrained to be nonnegative, this becomes a one-sided soft-thresholding step:

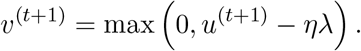

This step shrinks each weight by an amount proportional to *λ*. If a weight becomes small enough, it is set exactly to zero, which removes the corresponding gene.

Third, project the weights back into the allowed interval [0, 1]:

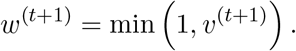

Equivalently, the full update can be written as

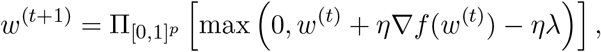

where 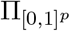 denotes projection onto the box [0, 1] .

Finally, the parameter *λ* controls the tradeoff between preserving spatial variation and selecting fewer genes. When *λ* is small, the sparsity penalty is weak, so more genes are selected. When *λ* is large, the penalty is stronger, so fewer genes remain. To tune *λ*, we evaluate a grid of candidate values. For each value of *λ*, the optimization produces weights *w*_*λ*_, selected genes

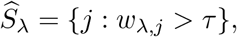

and the number of selected genes

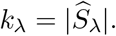

The preserved variation for that value of *λ* is

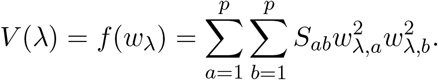

We then compare different values of *λ* using plots such as preserved variation versus *λ*, number of selected genes versus *λ*, and preserved variation versus number of selected genes. These plots show how much spatial co-expression variation is retained as the representation becomes sparser. A useful value of *λ* should select the set of genes that preserves a desired proportion of the spatial variation.

